# Distribution-Aware Federated Learning for Diabetes Prediction Using Tabular Clinical Data Under Non-IID and Class-Imbalanced Settings

**DOI:** 10.64898/2026.03.05.709751

**Authors:** Rohul Amin, Md. Mehedi Hasan Rana, Sumya Aktar

**Author notes:** Contributing authors. These authors contributed equally to this work.

## Abstract

Federated learning (FL) enables collaborative clinical model training without centralized data sharing, yet its deployment is hindered by statistical heterogeneity (non-IID data) and inherent class imbalance across healthcare institutions. Conventional aggregation strategies such as FedAvg and FedProx weight client updates solely by dataset size, ignoring class distributions and thereby biasing the global model toward the majority class. To address this, we propose Distribution-Aware Federated Learning (DA-FL), which introduces a minority-class amplification factor ***ϕ_k_*** computed as the ratio of a client’s local positive class rate to the global positive class rate. Combined with class-weighted cross-entropy loss at the client level, DA-FL forms a two-level correction mechanism that mitigates imbalance without additional data sharing. Experiments on the CDC BRFSS 2021 diabetes dataset (236,378 records across five simulated clients under three non-IID levels) show that DA-FL improves F1-Macro by 18.2% and G-Mean by 26.7% over FedAvg under moderate non-IID conditions, while achieving 31-fold greater F1-Macro stability across 30 communication rounds. These findings demonstrate that DA-FL is an effective and practically deployable solution for federated clinical prediction under realistic non-IID and class-imbalanced settings.

## 1 Introduction

The proliferation of electronic health records and wearable health monitoring devices has generated unprecedented volumes of patient data, creating significant opportunities for developing data-driven predictive models for chronic disease management [1]. Diabetes mellitus, affecting over 537 million adults globally, represents one of the most pressing public health challenges of the 21st century [2]. Early and accurate prediction of diabetes risk can facilitate timely clinical intervention and substantially reduce long-term complications. Machine learning and deep learning approaches have demonstrated considerable promise in clinical risk prediction tasks; however, their effectiveness is fundamentally contingent upon access to large, diverse, and representative training datasets.

In practice, patient data is distributed across geographically dispersed healthcare institutions hospitals, clinics, and community health centers each maintaining independent data repositories governed by stringent privacy regulations including the Health Insurance Portability and Accountability Act (HIPAA) and the General Data Protection Regulation (GDPR) [3]. These regulatory frameworks prohibit the centralization of patient data, rendering conventional centralized machine learning pipelines infeasible in real-world clinical deployments.

Federated Learning (FL), introduced by McMahan et al. (2017), offers a compelling solution to this challenge by enabling collaborative model training across distributed clients without requiring raw data to leave its source [4]. In FL, each participating client trains a local model on its private data and transmits only model parameters to a central server, which aggregates them into a global model. This paradigm preserves data privacy while enabling knowledge sharing across institutions.

However, the practical deployment of FL in clinical settings faces two critical and interrelated challenges that remain inadequately addressed in the existing literature. First, patient data distributions vary substantially across healthcare institutions, a phenomenon termed statistical heterogeneity or non-Independent and Identically Distributed (non-IID) data [5]. Different hospitals serve demographically distinct patient populations, employ different diagnostic equipment, and observe different disease prevalences, resulting in heterogeneous local data distributions [6]. Standard FL aggregation strategies, particularly FedAvg, exhibit significant performance degradation under non-IID conditions due to client model drift during local training [7]. While subsequent methods such as FedProx and FedNova have partially addressed this issue through modifications to local training objectives, they remain insufficient for highly heterogeneous clinical data.

Second, clinical datasets inherently exhibit severe class imbalance, diabetic patients constitute a minority of the general population, resulting in datasets where negative cases substantially outnumber positive cases [8]. In the CDC Behavioral Risk Factor Surveillance System (BRFSS) 2021 dataset employed in this study, diabetic cases represent only 14.2% of all records, yielding an approximate 6:1 class imbalance ratio [18]. Standard FL aggregation strategies aggregate client model weights proportionally by local dataset size, entirely disregarding local class distributions. Consequently, clients with few minority class samples contribute disproportionately to the global model, systematically biasing it toward the majority class. This results in poor sensitivity for the minority (diabetic) class, the clinically critical outcome, as reflected by degraded F1-Macro and G-Mean scores [7].

To address these dual challenges simultaneously, this paper proposes Distribution-Aware Federated Learning (DA-FL), a novel aggregation strategy that incorporates local class distribution information into the server-side model aggregation process. Specifically, DA-FL introduces a minority-class amplification factor *ϕ*_*k*_ for each client, computed as the ratio of the client’s local positive class rate to the global positive class rate, which modulates the client’s contribution to the global model aggregation. Clients with higher local minority class representation receive amplified aggregation weights, directly compensating for the federation-wide class imbalance. Additionally, DA-FL employs class-weighted cross-entropy loss during local training, providing a complementary two-level correction for class imbalance at both the local training and global aggregation stages.

The primary contributions of this work are as follows:

- We propose DA-FL, a distribution-aware federated aggregation strategy that incorporates a minority-class amplification factor into the server-side aggregation weight computation, addressing class imbalance at the federation level without modifying client data or sharing class distribution statistics beyond local positive rates.
- We demonstrate that DA-FL achieves substantially superior and more stable performance compared to FedAvg and FedProx under non-IID and class-imbalanced conditions, improving F1-Macro by 18.2% and G-Mean by 26.7% over FedAvg while exhibiting 31x greater F1 stability across communication rounds.
- We conduct comprehensive experiments on the large-scale CDC BRFSS 2021 diabetes dataset under three non-IID severity levels controlled by a Dirichlet distribution parameter, providing systematic evaluation of DA-FL’s robustness across varying degrees of data heterogeneity.
- We provide an open-source simulation framework using the Flower federated learning library, enabling reproducibility and serving as a benchmark for future research on imbalanced federated clinical prediction.

The remainder of this paper is organized as follows. Section 2 reviews related work on federated learning, non-IID FL methods, and class imbalance in federated settings. Section 3 presents the system model and the proposed DA-FL methodology. Section 4 describes the experimental setup and results. Section 5 discusses the findings and their clinical implications. Section 6 concludes the paper and outlines future research directions.

## 2 Related Work

### 2.1 Federated Learning Fundamentals

Federated Learning was formally introduced by McMahan et al. (2017) as a communication-efficient framework for training machine learning models across decentralized clients without centralizing raw data [4]. Their proposed algorithm, Federated Averaging (FedAvg), aggregates locally trained model weights at a central server using a weighted average proportional to each client’s local dataset size. While FedAvg demonstrated strong performance under idealized conditions, subsequent studies revealed substantial performance degradation when client data distributions are heterogeneous, a condition referred to as non-Independent and Identically Distributed (non-IID) data. The non-IID problem arises from two primary sources: label distribution skew, where different clients observe different class proportions, and feature distribution skew, where the statistical properties of input features vary across clients [5]. These challenges are particularly pronounced in real-world deployments where data is collected under varying conditions across distributed institutions.

### 2.2 Addressing Non-IID Data in Federated Learning

The non-IID challenge has motivated a substantial body of research proposing modifications to both local training objectives and global aggregation strategies. Meng et al. (2024) proposed FedProx, which augments the local training objective with a proximal term that penalizes client model parameters for deviating from the current global model [10]. This regularization reduces client drift during local training and improves convergence stability under heterogeneous data distributions. Wang et al. (2020) identified a related but distinct phenomenon termed objective inconsistency, wherein clients performing different numbers of local update steps contribute disproportionately to the global model [11]. Their proposed method, FedNova, normalizes local updates prior to aggregation to correct this inconsistency. Karimireddy et al. (2020) proposed SCAFFOLD, which employs control variates to correct for client drift at the gradient level [12]. While these methods address statistical heterogeneity from the perspective of local training dynamics, they retain size-proportional aggregation strategies that remain insensitive to local class distributions, a critical limitation in the presence of class imbalance.

Clustered federated learning approaches, such as those proposed by Sattler et al. (2021), group clients with similar data distributions into clusters and train separate models per cluster [13]. While effective for reducing distributional divergence within clusters, such approaches require additional communication overhead for cluster identification and may not generalize well when client distributions are continuously shifting. Personalized federated learning methods, including pFedMe [14] and FedPer [15], address heterogeneity by allowing clients to maintain personalized model components while sharing global representations. These methods prioritize per-client performance over global model quality and are therefore less applicable to scenarios requiring a single deployable global model for clinical inference.

### 2.3 Federated Learning for Healthcare Applications

The application of federated learning to healthcare has received considerable attention owing to the inherent sensitivity of patient data and the distributed nature of clinical data repositories. Rieke et al. (2020) provided a comprehensive overview of FL’s potential in digital health, identifying data heterogeneity and class imbalance as the two primary barriers to effective federated clinical model training [16]. A study demonstrated the feasibility of federated brain tumor segmentation across multiple institutional sites, showing that FL could approach the performance of centralized training under favorable conditions [17]. Specific to diabetes prediction, several studies have explored machine learning approaches using publicly available datasets. Dataset studies using the CDC BRFSS dataset have demonstrated the utility of ensemble methods and neural network architectures for diabetes risk stratification [18]. However, to the best of the authors’ knowledge, no prior work has investigated federated learning for diabetes prediction on the CDC BRFSS 2021 dataset under non-IID simulation, nor has any study proposed a distribution-aware aggregation strategy specifically designed to address the combined challenge of statistical heterogeneity and class imbalance in this context.

### 2.4 Class Imbalance in Federated Learning

Class imbalance in federated learning represents an underexplored but critical challenge, particularly in clinical applications where minority class samples correspond to clinically significant outcomes [19]. Duan et al. (2019) proposed Astraea, a self-balancing FL framework that addresses global class imbalance through data augmentation and mediator-based rescheduling of client training [20]. While effective, this approach requires clients to share class frequency information and introduces significant additional communication overhead. Sarkar et al. (2020) investigated the impact of non-IID and imbalanced data on FL convergence, demonstrating that standard aggregation strategies consistently underperform on minority class detection metrics including F1-Macro and G-Mean [21]. More recently, approaches combining oversampling techniques such as SMOTE with federated training have been proposed, though these require local data augmentation prior to federation and do not address the aggregation-level bias introduced by size-proportional weighting.

In contrast to existing approaches, the proposed DA-FL method addresses class imbalance directly at the aggregation level through a computationally lightweight minority-class amplification factor, without requiring data augmentation, additional communication rounds, or sharing of raw class frequency statistics beyond local positive rates returned as training metadata. This design preserves communication efficiency and privacy while providing meaningful correction for federation-wide class imbalance.

## 3 Methodology

### 3.1 System Model

Consider a federated learning system comprising one central server and *K* distributed clients, where *K* = 5 in this study. Each client *k* ∈ { 1, 2, …, *K* } holds a private local dataset

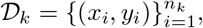

where *x*_*i*_ ∈ ℝ^*d*^ denotes the *d*-dimensional feature vector of the *i*-th sample and *y*_*i*_ ∈ { 0, 1 } denotes the corresponding binary class label. The total number of samples across all clients is

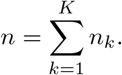

Raw data never leaves any client; only model parameters are transmitted between clients and the central server.

In this study, each client represents a simulated healthcare institution holding a subset of patient records from the CDC BRFSS 2021 dataset [22]. Client datasets are constructed to exhibit statistical heterogeneity through Dirichlet-based partitioning, reflecting a realistic scenario in which different institutions serve demographically distinct patient populations with varying diabetes prevalence rates. The overall system architecture of DA-FL System is given in Figure 1

**Fig. 1.**
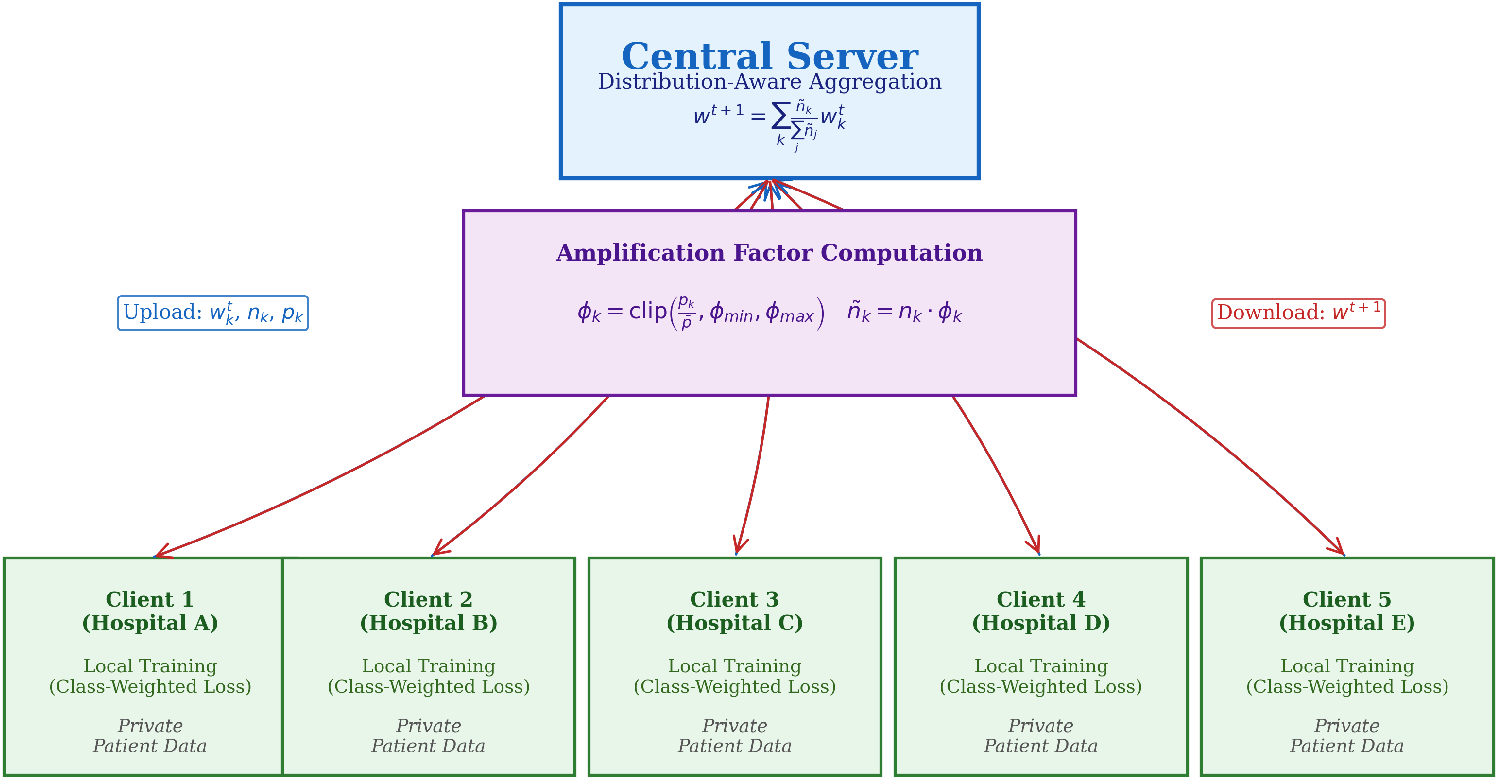
DA-FL System Architecture for Distribution-Aware Federated Learning for Clinical Diabetes Prediction

### 3.2 Problem Formulation

The global federated learning objective is to find model parameters *w*^∗^ that minimize the weighted aggregate of local loss functions:

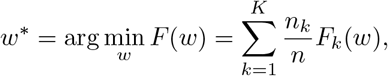

where *F*_*k*_(*w*) denotes the local objective function of client *k* and 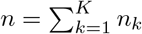.

Under standard FedAvg, the local objective is the binary cross-entropy loss:

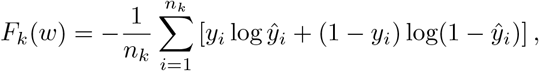

where *ŷ*_*i*_ denotes the predicted probability for sample *i*.

This formulation presents two critical limitations in the context of non-IID clinical data:

#### Aggregation insensitivity to class distribution

The aggregation weight 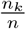 is determined solely by dataset size, without consideration of local class distributions. Clients with large datasets but very few minority-class samples can disproportionately bias the global model toward the majority class.

#### Local loss insensitivity to class imbalance

The standard cross-entropy loss treats all misclassifications uniformly, which causes local models to favor the majority class during training, particularly when the local positive rate is low.

DA-FL addresses both limitations through a two-level correction mechanism applied at (i) the local training stage and (ii) the global aggregation stage, respectively.

### 3.3 Local Training with Class-Weighted Loss

To address Limitation 2, each client *k* computes a local class weight *ω*_*k*_ based on its local class distribution:

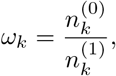

where 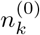 and 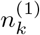 denote the number of negative and positive samples in client *k*’s local dataset, respectively.

The local training objective is then reformulated as a class-weighted binary cross-entropy loss:

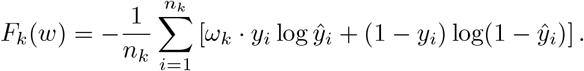

This formulation penalizes misclassification of minority-class samples by a factor of *ω*_*k*_ relative to majority-class misclassifications. Consequently, each client’s local model is encouraged to maintain sensitivity to the diabetic class irrespective of local class prevalence.

### 3.4 Distribution-Aware Global Aggregation

To address Limitation 1, DA-FL introduces a minority-class amplification factor *ϕ*_*k*_ for each client *k*, defined as

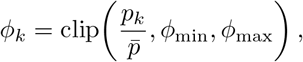

where 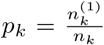 is client *k*’s local positive-class rate, 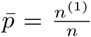 is the global positive-class rate across the federation, and clip(·, *ϕ*_min_, *ϕ*_max_) constrains *ϕ*_*k*_ to the interval [0.1, 5.0] to prevent any single client from dominating aggregation.

The combined aggregation weight for client *k* is then defined as

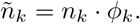

The global model at communication round *t* + 1 is computed as

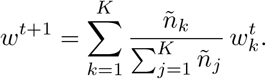

#### Intuition

Clients whose local positive rate exceeds the global rate 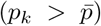 obtain *ϕ*_*k*_ > 1, thereby amplifying their contribution to the global model. Conversely, clients with substantially lower positive rates 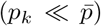 receive *ϕ*_*k*_ close to *ϕ*_min_ = 0.1, reducing their potentially biasing influence. This mechanism compensates for the under-representation of minority-class knowledge arising from uneven distribution of diabetic cases across clients.

#### Privacy preservation

The factor *ϕ*_*k*_ is computed using only the scalar local positive rate *p*_*k*_, which is transmitted as training metadata alongside model parameters. No raw samples, individual records, or detailed class histograms are shared with the server, thereby maintaining the privacy guarantees inherent to the federated learning framework.

### 3.5 The DA-FL Algorithm

The complete DA-FL procedure is formally described in Algorithm 1.

### 3.6 Model Architecture

Each client employs a Multilayer Perceptron (MLP) as the local model, comprising four fully connected layers with ReLU activation functions and dropout regularization. The architecture processes the 21-dimensional input feature vector through hidden layers of sizes 64, 128, and 64, before producing a single scalar output. Formally,

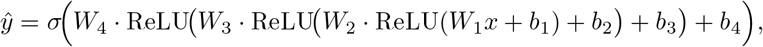

#### Algorithm 1

Distribution-Aware Federated Learning (DA-FL)

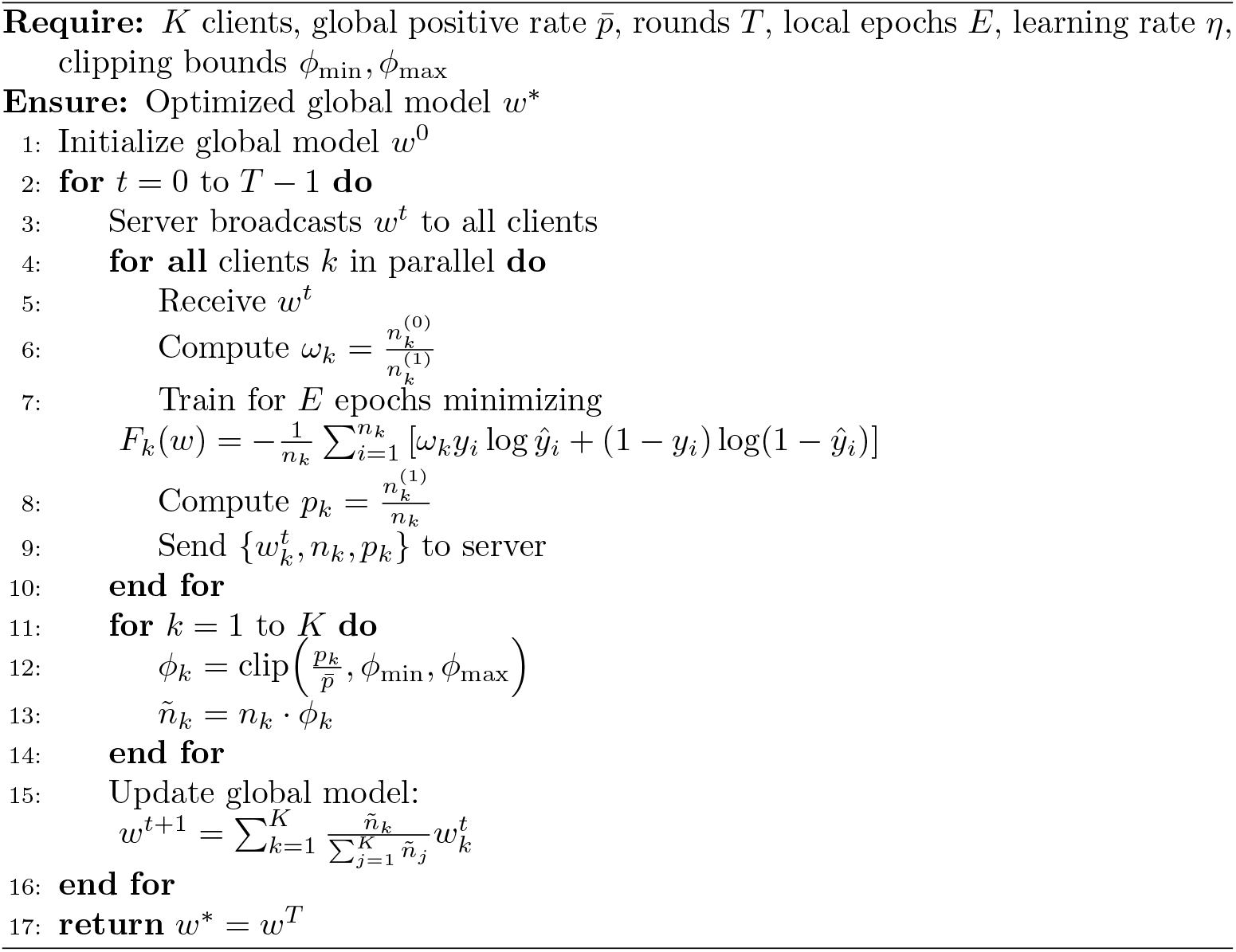

where *W*_*i*_ and *b*_*i*_ denote the weight matrix and bias vector of the *i*-th layer, respectively, and *σ*( · ) denotes the sigmoid activation applied at the output layer for probabilistic binary classification.

Dropout with rates of 0.3, 0.3, and 0.2 is applied after the first, second, and third hidden layers, respectively, to mitigate overfitting. The total number of trainable parameters is 18,049. Local training is performed using the Adam optimizer with learning rate *η* = 0.001.

The MLP architecture is selected due to its computational efficiency for tabular clinical data, compatibility with the Flower federated learning framework, and suitability for deployment on resource-constrained client devices, which is a practical requirement in real-world federated healthcare systems.

## 4 Experimental Setup

### 4.1 Dataset

Experiments are conducted using the CDC Behavioral Risk Factor Surveillance System (BRFSS) 2021 dataset, a large-scale publicly available health survey administered annually by the United States Centers for Disease Control and Prevention (CDC) [22]. The processed version employed in this study comprises 236,378 patient records and 21 clinical and demographic features, with a binary target variable indicating the presence or absence of diabetes mellitus. Features include physiological indicators (BMI, blood pressure, cholesterol), behavioral factors (physical activity, smoking, alcohol consumption), and socioeconomic attributes (education level, income, health-care access). The dataset exhibits a substantial class imbalance with 202,810 negative cases (85.8%) and 33,568 positive cases (14.2%), yielding an approximate 6:1 imbalance ratio. No missing values are present in the dataset. All continuous and ordinal features are standardized using z-score normalization prior to model training.

### 4.2 Federated Simulation Setup

All experiments are conducted using the Flower (flwr) federated learning simulation framework on a single computational instance, simulating K=5 distributed clients representing geographically dispersed healthcare institutions. The federation operates for T=30 global communication rounds. At each round, all K clients participate in local training and submit updated model weights to the server — corresponding to a client participation rate of 100%. Local training is performed for E=5 epochs per round using the Adam optimizer with learning rate *η*=0.001 and batch size of 64.

### 4.3 Non-IID Data Partitioning

Statistical heterogeneity across clients is simulated using Dirichlet distribution-based partitioning with concentration parameter *α*, following the established protocol of [23]. Three levels of non-IID severity are evaluated: *α* = 0.1 (extreme heterogeneity), *α* = 0.5 (moderate heterogeneity), and *α* = 1.0 (mild heterogeneity).

The resulting client-wise positive rates are summarized in Figure 2. At *α* = 0.1, individual clients exhibit positive rates ranging from 0.00% to 100.00%, representing the most challenging federated scenario in which some institutions observe exclusively a single class. At *α* = 0.5 and *α* = 1.0, distributions are progressively more balanced while retaining clinically realistic heterogeneity.

**Fig. 2.**
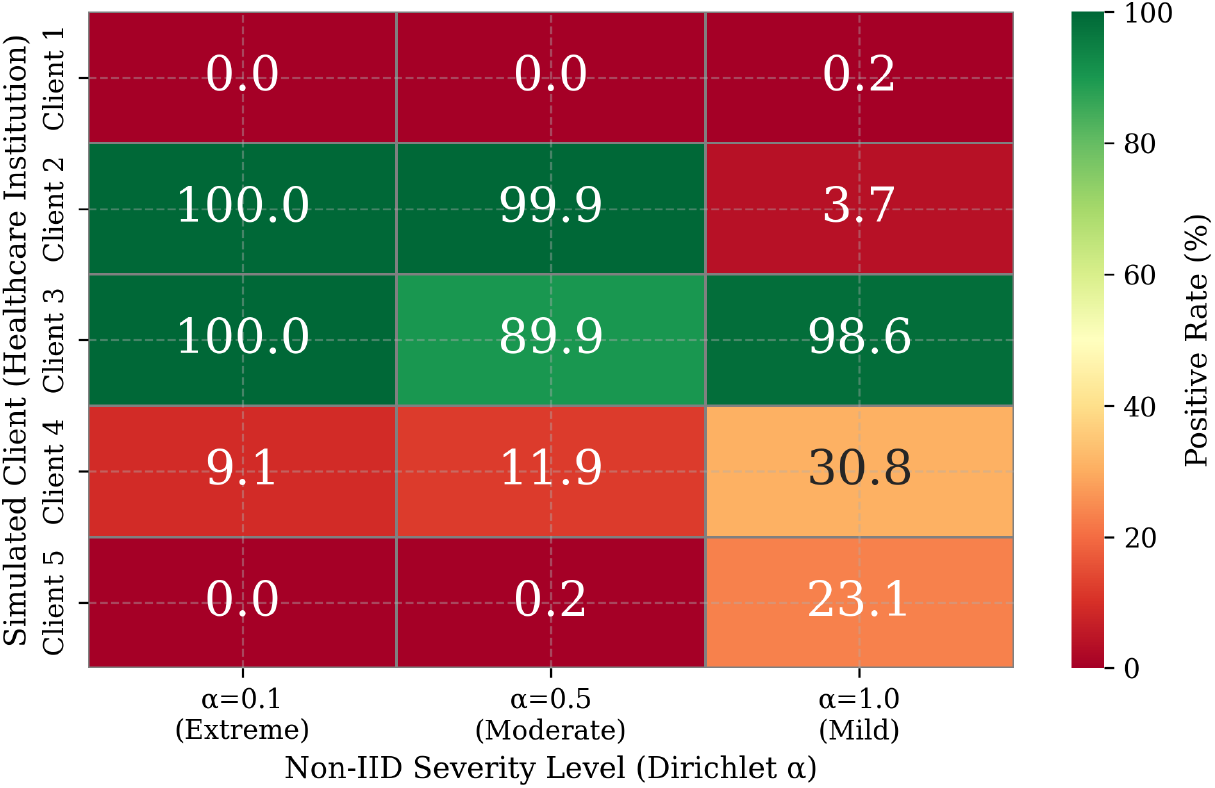
Client-wise Positive Rate (%) Across Non-IID Levels.

### 4.4 Baseline Methods

DA-FL is evaluated against the following baseline methods. Centralized Training trains all client data pooled at a single location without federation; this represents the theoretical performance upper bound and is included for reference only, as it violates data privacy constraints. Local only trains each client independently on its local data without any federation; this represents the lower bound and quantifies the benefit of collaborative training. FedAvg implements standard federated averaging with size-proportional aggregation weights and serves as the primary baseline. FedProx augments FedAvg with a proximal regularization term *µ* = 0.01 added to each client’s local loss function and serves as the non-IID-aware baseline.

All baseline methods employ identical local model architectures, optimizers, learning rates, batch sizes, and number of local epochs as DA-FL to ensure fair comparison. The sole differences between methods lie in the aggregation strategy and, for FedProx, the local loss function modification.

### 4.5 Evaluation Metrics

Given the clinical nature of the prediction task and the presence of significant class imbalance, evaluation relies on metrics that capture both overall and class-specific model performance.

#### Accuracy

Overall proportion of correctly classified samples. Reported for completeness but recognized as an inadequate sole metric under class imbalance.

#### Precision (Macro)

Average precision across both classes, treating each class equally regardless of size.

#### Recall (Macro)

Average recall across both classes. Particularly important clinically as it reflects the model’s sensitivity to the minority (diabetic) class.

#### F1-Macro

Harmonic mean of macro precision and recall. Primary metric for evaluating performance under class imbalance.

#### AUC-ROC

Area under the receiver operating characteristic curve. Measures the model’s discriminative ability independently of classification threshold.

#### G-Mean

Geometric mean of sensitivity and specificity, defined as

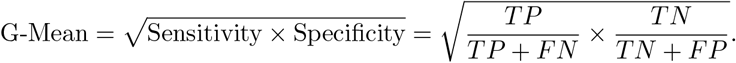

G-Mean is particularly suited to imbalanced classification as it simultaneously rewards high performance on both the majority and minority classes, penalizing models that achieve high accuracy by simply predicting the majority class.

All metrics are reported as weighted averages across clients, with client contributions weighted by local test set size.

### 4.6 Implementation Details

All experiments are implemented in Python 3.12 using PyTorch for local model training and the Flower (`flwr`) framework for federated simulation. Experiments are conducted on Google Colaboratory with CPU runtime. All random seeds are fixed at 42 to ensure reproducibility across data partitioning, model initialization, and training. The DA-FL amplification factor clipping bounds are set to *ϕ*_min_ = 0.1 and *ϕ*_max_ = 5.0 in all experiments. The complete source code is made publicly available to facilitate reproducibility.

## 5 Results

### 5.1 Overall Performance Comparison

Table 1 presents the performance of all methods under the primary experimental condition of *α* = 0.5 (moderate non-IID), representing the most clinically realistic federated scenario. The reported results correspond to evaluation metrics at Round 30, aggregated across all clients.

**Table 1.** Performance Comparison at *α* = 0.5 (Moderate Non-IID), Round 30.

| Method | Accuracy | Precision | Recall | F1-Macro | AUC-ROC | G-Mean |
| --- | --- | --- | --- | --- | --- | --- |
| FedAvg | 0.3342 | 0.5338 | 0.5997 | 0.2650 | 0.7773 | 0.4658 |
| FedProx | 0.5441 | 0.5288 | 0.7013 | 0.3955 | 0.7835 | 0.6631 |
| DA-FL (Proposed) | 0.6496 | 0.5328 | 0.7503 | 0.4471 | 0.7772 | 0.7329 |

DA-FL achieves the highest performance across all primary evaluation metrics. Relative to FedAvg, DA-FL improves Accuracy by 31.5%, Recall by 15.1%, F1-Macro by 18.2%, and G-Mean by 26.7%. Relative to FedProx, DA-FL improves Accuracy by 10.6%, Recall by 4.9%, F1-Macro by 5.2%, and G-Mean by 7.0%. AUC-ROC scores are comparable across all three methods (approximately 0.777–0.784), indicating that all methods maintain similar discriminative capacity while DA-FL substantially improves balanced classification performance.

The notably low Accuracy and F1-Macro of FedAvg (0.3342 and 0.2650, respectively) at Round 30 reflects model collapse toward the majority class under the combined pressures of non-IID data and class imbalance. In this scenario, the global model predominantly predicts the negative class, achieving high apparent specificity at the cost of near-zero sensitivity for diabetic cases. The G-Mean score of 0.4658 corroborates this interpretation, as a G-Mean below 0.5 indicates that the geometric balance between sensitivity and specificity falls below clinically acceptable thresholds.

DA-FL’s Recall of 0.7503 is particularly significant from a clinical perspective. In diabetes screening applications, false negatives, diabetic patients classified as healthy, carry substantially greater clinical risk than false positives. DA-FL correctly identifies 15.1% more diabetic patients than FedAvg at Round 30, representing a meaningful improvement in the clinical utility of the federated model.

### 5.2 Training Stability Analysis

Table 2 reports the mean, standard deviation, minimum, maximum, and range of F1-Macro, G-Mean, and AUC-ROC scores across all 30 communication rounds for each method under *α* = 0.5.

**Table 2.** Training Stability Analysis Across 30 Communication Rounds (*α* = 0.5)

| Method | Metric | Mean | Std | Min | Max | Range |
| --- | --- | --- | --- | --- | --- | --- |
| FedAvg | F1-Macro | 0.2904 | 0.1431 | 0.0882 | 0.4644 | 0.3762 |
| FedProx | F1-Macro | 0.2678 | 0.1309 | 0.0882 | 0.4469 | 0.3587 |
| DA-FL | F1-Macro | 0.4592 | 0.0046 | 0.4471 | 0.4668 | 0.0197 |
| FedAvg | G-Mean | 0.4520 | 0.2749 | 0.0000 | 0.7469 | 0.7469 |
| FedProx | G-Mean | 0.4185 | 0.2708 | 0.0000 | 0.7346 | 0.7346 |
| DA-FL | G-Mean | 0.7439 | 0.0342 | 0.5633 | 0.7617 | 0.1983 |
| FedAvg | AUC-ROC | 0.6775 | 0.1777 | 0.2625 | 0.8257 | 0.5632 |
| FedProx | AUC-ROC | 0.7046 | 0.1632 | 0.2336 | 0.8228 | 0.5893 |
| DA-FL | AUC-ROC | 0.7937 | 0.0106 | 0.7682 | 0.8075 | 0.0393 |

The stability results constitute the most compelling finding of this study. DA-FL exhibits an F1-Macro standard deviation of 0.0046, approximately 31 times lower than FedAvg (*σ* = 0.1431) and 28 times lower than FedProx (*σ* = 0.1309). Similarly, DA-FL’s AUC-ROC standard deviation of 0.0106 is 17 times lower than FedAvg (*σ* = 0.1777) and 15 times lower than FedProx (*σ* = 0.1632).

Both FedAvg and FedProx exhibit G-Mean scores of 0.000 in their worst rounds, indicating complete failure to detect minority-class samples at certain communication rounds. This represents a critical failure mode in clinical deployment, corresponding to rounds where the global model predicts exclusively negative outcomes for all patients. DA-FL’s worst G-Mean across all 30 rounds is 0.5633, indicating that even in its worst-performing round the model retains an acceptable sensitivity-specificity balance. This property is essential for reliable clinical federated deployment, where model behavior must be predictable and consistently safe across all training rounds. The AUC-ROC stability further underscores this finding. FedAvg’s AUC-ROC ranges from 0.2625 to 0.8257. A range of 0.5632 indicating near-random to strong discrimination across different rounds. DA-FL’s AUC-ROC range of 0.0393 represents a 14-fold improvement in discriminative stability. This consistency is essential for federated healthcare systems where model updates are deployed incrementally and unpredictable performance degradation between rounds poses patient safety risks.

### 5.3 Performance Across Non-IID Severity Levels

Table 3 and Figure 4 presents the Round 30 performance of all methods across three non-IID severity levels, providing a systematic evaluation of DA-FL’s robustness under varying degrees of statistical heterogeneity. Also, the convergence behavior of the FL methods under Non-IID and Class-Imbalanced condition is represented in Figure 3.

**Table 3.** Performance Comparison Across Non-IID Severity Levels (Round 30)

| $\alpha$ | Method | Accuracy | F1-Macro | AUC-ROC | G-Mean | Recall |
| --- | --- | --- | --- | --- | --- | --- |
| 0.1 (Extreme) | FedAvg | 0.7819 | 0.6036 | 0.7767 | 0.6996 | 0.7058 |
|  | FedProx | 0.8252 | 0.6316 | 0.7926 | 0.6866 | 0.7027 |
|  | DA-FL | 0.7731 | 0.5983 | 0.7777 | 0.7020 | 0.7067 |
| 0.5 (Moderate) | FedAvg | 0.3342 | 0.2650 | 0.7773 | 0.4658 | 0.5997 |
|  | FedProx | 0.5441 | 0.3955 | 0.7835 | 0.6631 | 0.7013 |
|  | DA-FL | 0.6496 | 0.4471 | 0.7772 | 0.7329 | 0.7503 |
| 1.0 (Mild) | FedAvg | 0.4385 | 0.3408 | 0.7756 | 0.5748 | 0.6429 |
|  | FedProx | 0.4695 | 0.3617 | 0.7702 | 0.5996 | 0.6550 |
|  | DA-FL | 0.6764 | 0.4697 | 0.7881 | 0.7153 | 0.7191 |

**Fig. 3.**
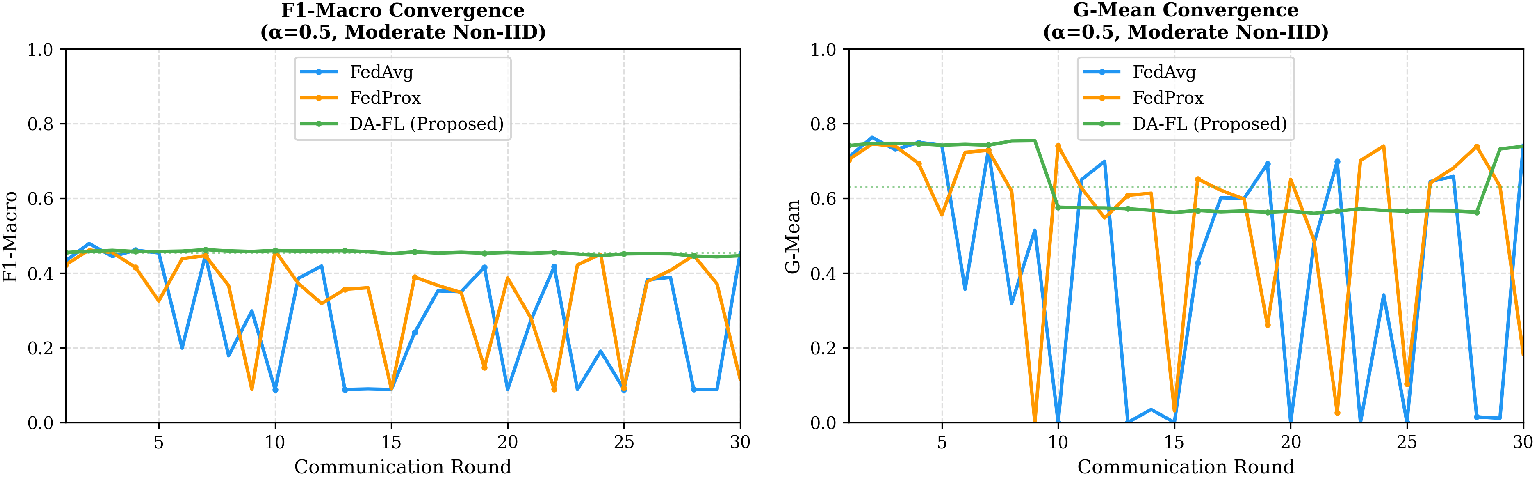
Convergence Behavior of FL Methods Under Non-IID and Class-Imbalanced Conditions(K=5 Clients, T=30 Rounds).

**Fig. 4.**
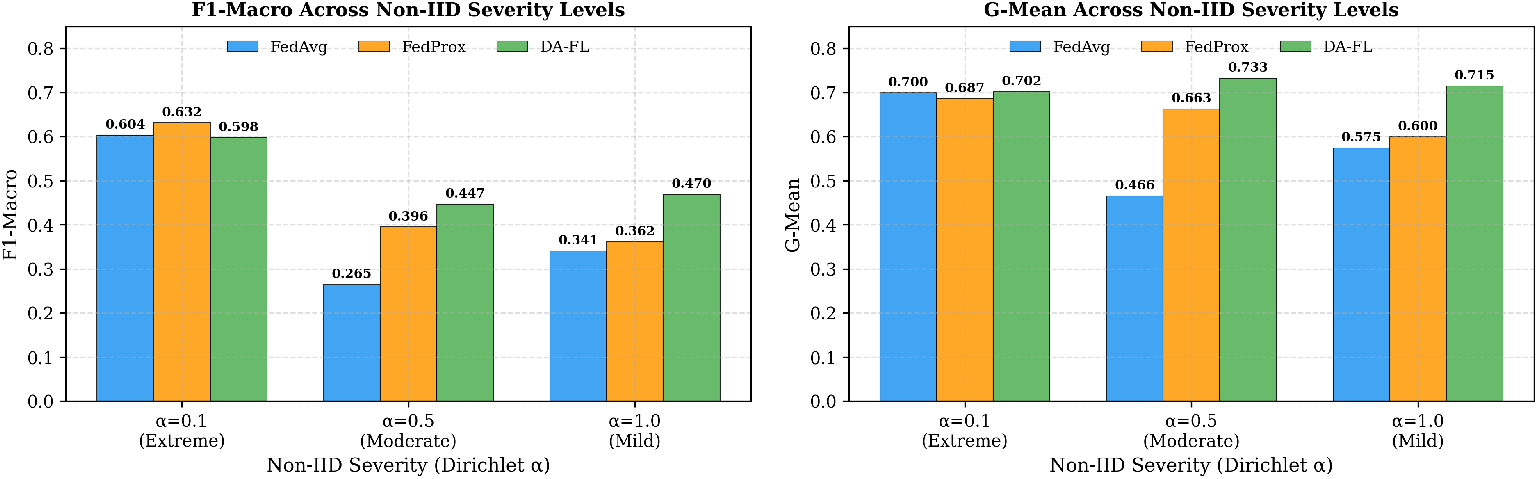
Performance Comparison Across Non-IID Severity Levels.

DA-FL demonstrates consistent superiority under moderate (*α* = 0.5) and mild (*α* = 1.0) non-IID conditions across all primary evaluation metrics. Under mild non-IID conditions (*α* = 1.0), DA-FL achieves an F1-Macro of 0.4697 and a G-Mean of 0.7153, improving over FedAvg by 12.9% and 14.1% respectively, and over FedProx by 10.8% and 11.6% respectively.

Under extreme non-IID conditions (*α* = 0.1), DA-FL demonstrates competitive performance on clinically critical metrics, achieving the highest G-Mean (0.7020) and Recall (0.7067) among all methods, while FedProx achieves marginally superior F1-Macro (0.6316 versus 0.5983) and Accuracy (0.8252 versus 0.7731). This relative underperformance of DA-FL on F1-Macro at *α* = 0.1 is attributable to the polarization of the amplification factor *ϕ*_*k*_ under extreme partitioning conditions, where clients with entirely single-class data receive maximally clipped *ϕ*_*k*_ values of either 5.0 or 0.1. In such cases, the amplification mechanism concentrates aggregation weight onto a small number of clients, reducing the diversity of the global model update. Notably, DA-FL retains its advantage on G-Mean even at *α* = 0.1, indicating that minority class sensitivity, the most clinically critical property, is preserved across all heterogeneity levels.

These results suggest that DA-FL is most effective in federated healthcare scenarios characterized by moderate to mild institutional heterogeneity, conditions that are arguably the most representative of real-world multi-institutional deployments, where complete single-class data partitioning is unlikely in sufficiently large patient populations.

### 5.4 Analysis of Distribution-Aware Aggregation Weights

Table 4 presents the aggregation weights assigned to each client by DA-FL at representative communication rounds, illustrating the mechanism’s behavior under the *α* = 0.5 partition.

**Table 4.** DA-FL Aggregation Weights at Representative Rounds (*α* = 0.5)

| Client | Samples | Positive Rate | $\phi_k$ | Combined Weight | Weight % |
| --- | --- | --- | --- | --- | --- |
| Client 1 | 42,328 | 0.01% | 0.10 | 4,232 | 2.5% |
| Client 2 | 22,227 | 99.91% | 5.00 | 11,135 | 6.7% |
| Client 3 | 22,948 | 89.90% | 5.00 | 114,740 | 68.9% |
| Client 4 | 31,660 | 11.95% | 0.84 | 26,631 | 16.0% |
| Client 5 | 89,937 | 0.24% | 0.10 | 8,993 | 5.4% |

Under standard FedAvg, Client 5, with 89,937 samples but only 0.24% positive rate, would receive the highest aggregation weight (38.0% by dataset size alone), disproportionately biasing the global model toward the majority class. DA-FL reduces Client 5’s effective weight to 5.4% while amplifying Client 3’s contribution to 68.9%, reflecting its substantially higher minority class representation (89.90% positive rate). This redistribution of aggregation influence directly explains DA-FL’s improved minority class sensitivity and G-Mean performance.

The experimental results collectively demonstrate that distribution-aware aggregation constitutes an effective and practical approach to addressing class imbalance in federated clinical prediction. The proposed *ϕ*_*k*_ mechanism operates entirely at the server side, requires no modification to client data or local training procedures beyond the return of a single scalar positive rate value, and introduces negligible computational overhead, with additional computation per aggregation round of *O*(*K*), where *K* is the number of clients.

The two-level imbalance correction employed by DA-FL, comprising class-weighted local loss and distribution-aware aggregation, addresses the problem from complementary perspectives. Class-weighted loss ensures that each client’s local model maintains minority class sensitivity regardless of local prevalence. Distribution-aware aggregation ensures that the global model disproportionately inherits knowledge from clients with meaningful minority class representation, correcting the systematic bias introduced by size-proportional aggregation under heterogeneous distributions.

From a practical deployment perspective, the stability properties of DA-FL are arguably more important than peak performance metrics. In real federated healthcare systems, model updates are deployed incrementally to clinical decision support tools, and unpredictable oscillation in model performance between rounds, as exhibited by FedAvg and FedProx, poses patient safety risks that are unacceptable in clinical environments. DA-FL’s consistent maintenance of G-Mean above 0.56 across all 30 rounds provides reliability guarantees that neither FedAvg nor FedProx can offer under the evaluated conditions.

### 5.5 Limitations

Several limitations of the present study warrant acknowledgment. First, client heterogeneity in this study is simulated through Dirichlet partitioning of a single centralized dataset, which approximates but does not fully replicate the feature distribution shifts arising from genuine institutional differences in data collection protocols, patient demographics, and clinical workflows. Second, the proposed amplification factor *ϕ*_*k*_ exhibits reduced effectiveness under extreme non-IID conditions (*α* = 0.1), where client distributions are entirely polarized, suggesting that adaptive clipping strategies may further improve performance in such scenarios. Third, experiments are conducted with *K* = 5 clients, a relatively small federation, and the scalability of DA-FL to larger federations with *K* = 20 or *K* = 50 clients represents an important direction for future investigation. Fourth, the MLP architecture employed, while appropriate for tabular clinical data, may not generalize to federated scenarios involving imaging or multimodal clinical data without architectural adaptation.

## 6 Conclusion

This paper presented Distribution-Aware Federated Learning (DA-FL), a novel aggregation strategy addressing statistical heterogeneity and class imbalance in federated clinical prediction. DA-FL introduces a minority-class amplification factor *ϕ*_*k*_ at the server-side aggregation stage, modulating each client’s contribution based on its local positive class rate relative to the global rate. Combined with class-weighted cross-entropy loss locally, DA-FL provides a two-level correction mechanism that compensates for majority-class bias under non-IID and class-imbalanced conditions.

Experiments on the CDC BRFSS 2021 diabetes dataset with five simulated clients across three non-IID levels show that DA-FL consistently outperforms FedAvg and FedProx on clinically relevant metrics. Under moderate non-IID (*α* = 0.5), DA-FL improves F1-Macro by 18.2% and G-Mean by 26.7% over FedAvg while maintaining comparable AUC-ROC. DA-FL also achieves dramatically superior training stability, with 31-fold and 14-fold reductions in F1-Macro and AUC-ROC variance, respectively, across 30 rounds. Under extreme non-IID conditions (*α* = 0.1), DA-FL retains competitive performance on G-Mean and Recall, delineating its operational envelope.

DA-FL is architecture- and dataset-agnostic, introduces minimal computational overhead (*O*(*K*) per aggregation round), and is suitable for communication- and compute-constrained federated environments. Future work includes extending *ϕ*_*k*_ to multi-class classification, integrating adaptive clipping for extreme distributions, evaluating compatibility with differential privacy, scaling to larger federations with partial client participation, and adapting the method to multimodal clinical data.

## Declarations

### Ethics approval and consent to participate

Not applicable.

### Consent for publication

Not applicable.

### Data availability

The dataset used in this study is publicly available at the CDC BRFSS 2021 repository: https://www.cdc.gov/brfss/annual_data/annual_2021.html.

### Materials availability

Not applicable.

### Code availability

The code used for experiments is publicly available at: https://github.com/labidz/FedLearn/blob/main/fedleanr(2).py.

### Author contribution

Rohul Amin conceived the study, designed the methodology, performed the main experiments, analyzed the results, and drafted the manuscript. Md. Mehedi Hasan Rana contributed to the preprocessing of the dataset, assisted in implementing parts of the experimental setup, and participated in reviewing and editing the manuscript. Sumya Aktar contributed to feature selection, assisted in interpreting the results, and helped refine the manuscript. All authors read, revised, and approved the final version of the manuscript, and agree to be accountable for all aspects of the work.

